# Concealed identity information detection with pupillometry in rapid serial visual presentation

**DOI:** 10.1101/2021.06.18.448944

**Authors:** Ivory Y. Chen, Aytaç Karabay, Sebastiaan Mathot, Howard Bowman, Elkan G. Akyürek

## Abstract

The concealed information test (CIT) relies on bodily reactions to stimuli that are hidden in mind. However, people can use countermeasures, such as purposely focusing on irrelevant things, to confound the CIT. A new method designed to prevent countermeasures uses rapid serial visual presentation (RSVP) to present stimuli on the fringe of awareness. Previous studies that used RSVP in combination with electroencephalography (EEG) showed that participants exhibit a clear reaction to their real first name, even when they try to prevent such a reaction (i.e., when their name is concealed information). Since EEG is not easily applicable outside the laboratory, we investigated here whether pupil size, which is easier to measure, can also be used to detect concealed identity information. In our first study, participants adopted a fake name, and searched for this name in an RSVP task, while their pupil sizes were recorded. Apart from this fake name, their real name and a control name also appeared in the task. We found pupil dilation in response to the task-irrelevant real name, as compared to control names. However, while most participants showed this effect qualitatively, it was not statistically significant for most participants individually. In a second study, we preregistered the proof-of-concept methodology and replicated the original findings. Taken together, our results show that the current RSVP task with pupillometry can detect concealed identity information at a group level. Further development of the method is needed to create a valid and reliable concealed identity information detector at the individual level.

## 1 Introduction

Concealed information is something you hide from others in your mind. Concealed information can be anything, from your opinion on the clothing habits of your colleagues, to more serious crime-related information, for instance about a tool used in a crime, a particular date when a crime was or is going to be carried out, or identity information such as the name of a victim or an accomplice (Suchotzki & Gamer, 2018). To find reliable ways to detect such concealed crime-relevant information has long been a major goal of forensic scientists. With a reliable and valid method of detecting concealed information, more crimes could be solved and guilt as well as innocence could be more readily established.

The concealed information test (CIT) is a method that has been developed for this purpose. It has a high validity to detect concealed information, and it has been gradually improved in the past years (Ben-Shakhar & Elaad, 2003; Meijer et al., 2014; Verschuere & Kleinberg, 2016; Volz et al., 2018). The CIT was originally created by Lykken (1959) to test whether participants have crime-relevant knowledge. Generally, in the CIT, testers show participants crime-relevant stimuli (i.e., stimuli related to information that only the perpetrator has), and some neutral alternatives. The CIT rests on the assumption that implicit responses will be evoked by crime-relevant stimuli if participants already have that “guilty” knowledge. Thus, responses to the crime-relevant stimuli are compared to the responses to the neutral alternatives to assess whether the participant indeed has crime-relevant knowledge (Ben-Shakhar et al., 2011).

Several different response measures in the CIT have been developed with varying levels of success. These measures include autonomic-nervous-system responses that indicate levels of arousal, such as heart rate, respiration, and electrodermal activity (Kleiner, 2002; Rosenfeld et al., 2007), as well as eye movements, such as fixations, saccades, blinks, and pupil responses (Gamer & Pertzov, 2018; Janisse & Bradley, 1980). In recent years, neuroimaging techniques, such as functional magnetic resonance imaging (fMRI), and stimulus-evoked brain potentials from electroencephalography (EEG), have become popular tools to record the reaction of the brain to stimuli in the CIT (Gamer, 2014; Ganis, 2014; Hu et al., 2011; Mameli et al., 2010; Zeki et al., 2004; but see also Furedy, 2009, for a note on applicability).

Nevertheless, despite extensive research efforts, the CIT cannot always reliably detect concealed information (Matsuda et al., 2012; Meijer et al., 2016). Its validity has been questioned because examinees can purposefully use physical and mental countermeasures to obscure the difference between responses to relevant and neutral alternatives (Ben-Shakhar, 2011; Peth et al., 2016). For example, they can bite their tongues to inflict pain or recall exciting memories when neutral alternatives are presented, and thereby confound the measure (Mertens & Allen, 2008; Rosenfeld et al., 2004). The usefulness of any test that is only reliable with fully compliant examinees is obviously limited, and it is therefore important to find solutions to defeat countermeasures.

A new method, based on rapid serial visual presentation (RSVP), has recently been developed with the potential to eliminate this problem. In RSVP, a series of stimuli are presented sequentially in the same location with each stimulus visible only for about 100 milliseconds (Broadbent & Broadbent, 1987). This quick presentation on the fringe of awareness virtually eliminates the possibility of using countermeasures, because participants do not have enough time to exert top-down control over their responses to the stimuli. Bowman et al. (2013) first developed the RSVP paradigm for concealed identity information detection. The researchers measured EEG during a fake-name search task, and found that participants’ real names triggered significant P3 potentials compared to control names, even when participants were explicitly instructed to hide their responses to their real name in various ways (Bowman et al., 2014). The results of Bowman and colleagues suggest that the RSVP paradigm reveals concealed identity information, is robust to countermeasures, and is therefore potentially more effective than the slower methods of presentation used to date.

Although the EEG results obtained by Bowman and colleagues (2013, 2014) with their RSVP task are promising, their method is not yet suitable for widespread application. Consider, for instance, that EEG facilities are not commonplace outside of university laboratories. Also with the CIT, it has been noted that new measures should be added to increase the usefulness of the CIT as a source of evidence in court (Matsuda et al., 2012). It is thus important to assess whether the RSVP method is also effective with simpler measures that are more readily available in practice. Eye movements, which can be recorded easily and unobtrusively, may be one such measure.

Eye movements reflect various cognitive processes that are also relevant for the CIT, such as task-directed cognitive control and visual attention. Several studies found that the rate of spontaneous eye blinks is positively correlated with dopamine levels at a neurobiological level, and with task-directed behavior at a behavioral level (Eckstein et al., 2017). In addition, the rate of microsaccades is controlled by the superior colliculus (SC), which is also involved in voluntary-saccade target selection (Hafed et al., 2009).

Eye movements have also been shown to be an effective independent measure in CIT. In Millen et al. (2017)’s study, some faces that were new, recently learned, or highly familiar to participants were presented sequentially and participants were asked to classify these faces based on familiarity. Regardless of whether participants were instructed to conceal their familiarity or not, highly familiar faces triggered fewer fixations to fewer regions, together with longer fixation durations. Similar results were obtained in other studies (Althoff et al., 1999; Heisz & Shore, 2008; Peth et al., 2013, 2016; Ryan et al., 2000, 2007). In addition, both barely visible familiar faces and stimuli from pre-learned text and videos produced earlier and longer inhibition of microsaccades as well as inhibition of eye blinks. Based on these measures, ‘terrorists’ could be distinguished from ‘innocents’ (Rosenzweig & Bonneh, 2019, 2020). Taken together, these results suggest that eye movements, for which no sensors or electrodes need to be attached to a suspect, and for which only a sensitive video camera (i.e., an eye tracker) is needed, might also be an efficient measure for RSVP-based detection of concealed information.

In this study, we focus on a particular kind of eye movement, namely the pupil response. Pupil responses are a promising measure for RSVP-based concealed information testing. First, pupil dilation and the P3 component of the event-related potential both reflect phasic responses in the locus coeruleus-norepinephrine (LC-NE) system. According to Nieuwenhuis et al. (2011), a motivationally significant stimulus will evoke a dilation of the pupils as well as a P3. These two reactions are tightly linked to the activation of the LC-NE system (Koss, 1986; Murphy et al., 2011; Nieuwenhuis et al., 2005; Samuels & Szabadi, 2008). Processing task-relevant events will activate LC-NE’s phasic response, followed by a pupil dilation and the P3 (Aston-Jones & Cohen, 2005). Since the P3 has proved to be an effective measure in RSVP-based CIT studies, it is reasonable to suppose that pupil size will also be useful as a measure in the RSVP-based CIT.

Second, pupil dilation is capable of showing two different cognitive control and attentional processes to task-relevant and task-irrelevant information in the CIT. Pupil dilation has been found to reflect the degree of cognitive control and attention required for responding to task-relevant information (i.e., target stimuli), while inhibiting irrelevant distractors (Cohen et al., 2015; Querino et al., 2015; Rondeel et al., 2015; van der Wel & van Steenbergen, 2018). Similarly, a critical concealed-information stimulus, despite being task-irrelevant, will also attract attention and engage cognitive control processes. For instance, it has been shown that the pupils also dilate when attention is allocated to new and salient stimuli that are task-irrelevant (Gilzenrat et al., 2010).

In addition, like other eye movements, pupil size has already been used in CIT studies and found to be an effective measure of concealed information. Lubow & Fein (1996) trained participants to be either guilty or innocent in a mock-crime scenario, and then showed them some photographs of crime-relevant items (e.g., a green identification card or a face of a criminal), together with some crime-irrelevant items. 50-70% of the guilty participants and 100% of the innocent participants were correctly detected through the difference between pupil sizes to the crime-relevant and crime-irrelevant items. Another study also adopted pupil size as an indicator of concealed mock-crime-related knowledge (Seymour et al., 2013). With a hit rate of 83% and zero false-alarms, the authors were able to distinguish guilty from innocent participants with 92% accuracy.

Finally, even though the pupil response is slow, which at first glance might raise questions as to its effectiveness in an RSVP task, Wierda et al. (2012) showed that pupil dilation is able to reflect attention allocation and cognitive processing in RSVP. In their study, they used an attentional blink task of two target letters within a sequential stream of digits as distractors presented in an RSVP. Using a pupil-dilation deconvolution method, the occurrence and timing of attentional processes, associated with target detection, were clearly tracked.

To sum up, the findings reviewed above support the idea that pupil size could be an effective measure in RSVP-based CIT. The aim of the current studies was thus to test the ability of the RSVP method in combination with a measure of pupil dilation to detect concealed identity information.

## 2 Experiment 1

For the aforementioned purpose, we used the design of the study by Bowman and colleagues, with some minor changes for pupil data recording, and using pupillometry instead of EEG. Each trial consisted of an RSVP stream, in which either a fake name, the participants’ real name or a randomly selected control name appeared. The participants were asked to search for the fake name and to ignore their real name. Their pupil sizes were recorded during the task, and we tested whether their pupil responses to the real and control names differed reliably. The presence of such a difference would be indicative of the ability of the RSVP-pupil method to detect concealed identity information.

### 2.1 Method

#### 2.1.1 Participants

Thirty-one participants took part in the experiment. All of them were first-year undergraduate students at the University of Groningen in the age group of 18-24 years (Mean: 19.29 years); there were 26 females and 5 males. All participants were native Dutch speakers. 26 participants were right-handed and 5 were left-handed. Participants had normal (uncorrected) vision. During the experiment, participants did not wear glasses, eye contacts or eye make-up. The study was conducted in accordance with the World Medical Association Declaration of Helsinki (2013) and approved by the ethical committee of the Psychology Department of the University of Groningen (approval number: PSY-18167-SP). Written informed consent was obtained prior to participation. Written and oral debriefing was provided after participation. Participants received course credits as compensation.

#### 2.1.2 Apparatus and Stimuli

Participants were seated with their head on a chin rest with an adjustable height at a distance of approximate 60 cm from a 27’’ LCD Iiyama PL2773H monitor with a display resolution of 1920×1080 pixels and a refresh rate of 100 Hz. On this monitor, stimuli were presented with OpenSesame 3.2.8 (Mathôt et al., 2012) running on Windows 10 Enterprise. Pupil size was recorded in arbitrary units by an EyeLink eye tracker (SR Research) during each trial using PyGaze (Dalmaijer et al., 2014).

We created a set of names for the experiment based on a database from the Meertens Institute for Dutch language and culture research (Https://www.Meertens.Knaw.Nl/Nvb/Topnamen/Land/Nederland). We first selected the 100 top Dutch names of each year from 1975 to 2014. Next, we excluded names consisting of more than 10 letters. This resulted in a set of 533 names with 281 female and 252 male names. From this name set, prior subsets of 15 possible names were selected randomly for each participant. A fake name and a control name were both selected from the unfamiliar names in the prior name subsets. Additionally, distractor names in each trial were selected pseudo-randomly from the set of 533 names: names with more than two identical consecutive letters were not allowed to be next to each other in one sequence. For example, ‘Dani’ and ‘Daniel’ had four identical consecutive letters; therefore, these two names could not be shown to participants directly after each other in one sequence. The distractor names were only used to form each name sequence and their presentation frequency was far less than the frequency of the fake, real and control names, which we were primarily concerned with.

We padded names on both sides with ‘+’ and ‘#’ characters so that the resulting string always consisted of 11 characters, as illustrated in Figure 1. Name stimuli were light grey (75% white; RGB: 190, 190, 190), 48 point size, sans serif characters presented on a dark (RGB: 40, 40, 40) background. All the names were presented in the center of the screen. The visual angle for each name was 2.03° in height and from 8.88°-12.25° in width, with some variation because some letters are wider than others. Fixation dots were light grey (75% white; RGB: 190, 190, 190), and rendered in 48 point.

**Figure 1.**
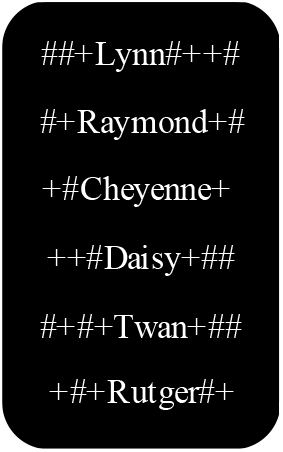
Examples of stimuli. List of example names used as stimuli. Names are padded in order to keep letters in the center positions.

#### 2.1.3 Procedure

Prior to the start of the experiment, participants were presented with a subset of 15 possible female or male names matching their own gender from the name set. They were asked to indicate all the names of people they knew; these names were removed from the set of possible names to avoid confounds due to the familiarity of such names. After that, participants chose one of the remaining names as their target name for detection during the experiment; we refer to this as their “fake” name. If a participant removed all 15 names, a second round of 15 names would be shown until a fake name was selected. The primary task for the participants was to monitor the RSVP streams for the presence of this fake name. A single “control” name would also be selected from the remaining names after the fake name was selected, and their own “real” name was also added to the experiment. The pupil sizes of participants when they saw these three different critical names were used for later analyses.

Once a fake name and a control name were selected, the experiment started. As shown in Figure 2, each trial started with a drift-correction procedure (i.e., a one-point recalibration) followed by a fixation dot presented for 1000 milliseconds in order to establish a baseline pupil size. Then a stream of 11 names were displayed for 100 milliseconds each in a sequence. A dashed line (----------) or series of equal signs (=======) was presented for 100 milliseconds after the sequence. Participants were required to report this later as a secondary task that served to check whether their attention had remained on the stimulus presentation area throughout the stream. At the end of each trial, a fixation dot was shown again for 2000 milliseconds, to allow capture of the full pupil response, which continues for some time after stimulus presentation. Participants were asked to try not to blink from the appearance of the first fixation dot to the disappearance of the second fixation dot.

**Figure 2.**
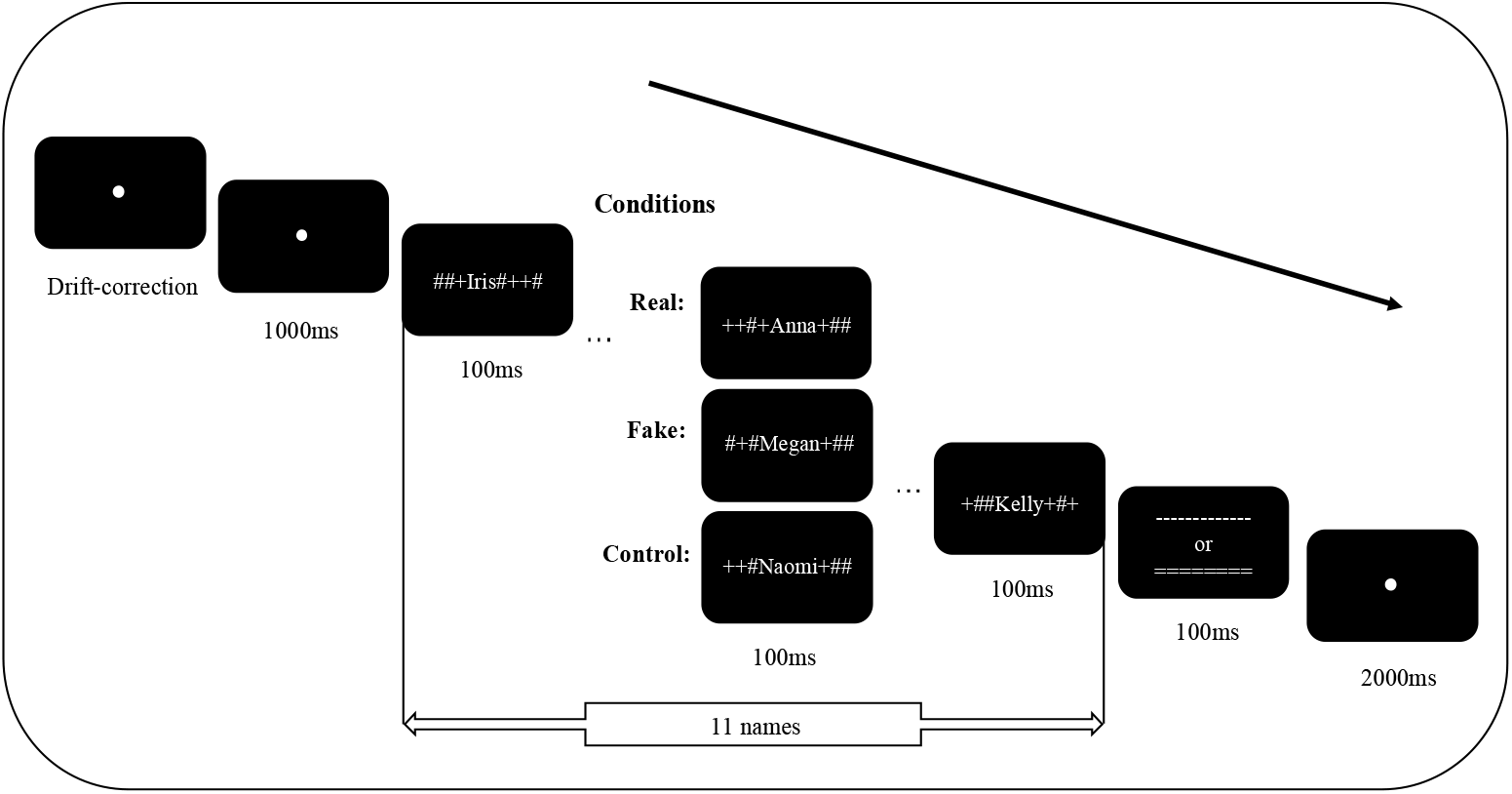
Trial sequence. In this example, Anna is the participant’s *real* name; Megan is the *fake* name chosen by the participant; and Naomi was selected as the *control* (irrelevant) name.

In the sequence, 10 of 11 names were distractors. One critical name, at a random position between 5 (earliest) and 9 (latest) in the sequence, defined three different conditions: this name was the participants’ real name (to which no behavioral response should be made), the fake name (the name that participants had selected previously and were instructed to respond to), or a control name (a name randomly selected from unfamiliar names of the previous sets with 15 names each, serving as a baseline that was matched in presentation frequency to the real and fake names, and which also did not require a response). Each condition was represented in 60 out of 180 trials separately.

Participants were instructed to ignore their real name and other irrelevant names, but pay attention to the fake name and the symbols at the end of each trial. After each presentation, participants were asked to answer two questions: 1. Did you see your (fake) name? 2. Did you see ---------- or =======? Participants answered the two questions by pressing the ‘F’ or ‘J’ keys on a standard QWERTY keyboard. Whether ‘F’ was for ‘Yes’ or ‘No’ was counter-balanced based on participant number parity to balance the response mapping between participants.

There were two sessions in the whole experiment: a practice session and an experimental session. The practice session consisted of 20 trials, during which participants received trial-based feedback (a smiley face for correct answers and a frowney face for wrong answers) to indicate whether they had responded correctly or not to the first and second question. The experimental session consisted of 180 trials divided into 6 blocks with a break between each pair of blocks. Conditions (real, fake, control) were randomly mixed within blocks. All participants took part in all three conditions, and the practice session, and thus completed 200 trials in total.

### 2.2 Data processing and analysis

All data and the analysis scripts are publicly accessible on the Open Science Framework (Https://osf.io/9fkpm/).

We first analyzed how accurately participants responded to question 1 (did you see your (fake) name?) and question 2 (did you see ---------- or =======?). The response accuracy for question 1 reflects how well participants were able to detect their fake names, and therefore whether the difficulty of the RSVP task was reasonable in the sense that it did not show a floor or ceiling effect. The accuracy for question 2 reflects to what extent participants maintained their attention on the RSVP stream throughout the trial, also after the appearance of the critical name.

On each trial, 800 samples of pupil-size data were obtained, given that the EyeLink eye tracker sampled at 250 Hz and a whole RSVP trial was 3200ms from the start of a name sequence to the end of the following fixation. We down-sampled the signal to 25 Hz, leaving 80 samples per trial for each participant. Then we baselined the pupil sizes by subtracting the mean of the first three samples (baseline) in the pupil traces for each trial separately. We calculated z scores of the baselines for each trial and excluded the trials that had a z score larger than 2 or less than −2 (Mathôt & Vilotijević, 2022). After baselining, we locked the pupil sizes to the onsets of the critical name positions, such that timepoint 0 corresponded to the onset of the name regardless of at which position in the stream the name was presented. Finally, because of the initial pupil constriction that occurs generically after the onset of stimuli (Mathôt, 2018), we deleted the first 200ms of data, leaving 2200 ms (55 samples) for each participant per trial.

We ran a sample-by-sample linear mixed effects analysis on the group level to check for possible effects on pupil size of the fake and real names, compared to the control names. More specifically, for each 40 ms sample separately we conducted a linear mixed effects analysis with baseline-corrected pupil size as dependent measure, condition as fixed effect (with control as reference value), and by-participant random intercepts and slopes, using the lme4 and lmerTest packages for R. For sample-by-sample analyses, we considered an effect reliable if *p* < .05 for at least 200 ms (5 samples).

Our predictions were twofold. First, if pupil size in the fake condition is significantly larger than in the control condition, then this indicates that the fake names (the task-relevant stimuli) elicited a reaction, and that this (presumably attentional) reaction can be detected by pupillometry in RSVP at the group level, as would be expected from prior research. Second, if pupil size in the real condition is significantly larger than in the control condition, then this indicates that the real name also elicited a detectable reaction, despite this name being task-irrelevant. This second effect is what we were primarily interested in, because it would indicate that pupil size is useful as a tool for the RSVP-based concealed-information detection.

In addition, we tested whether pupil size was modulated by learning, fatigue, or habituation over the course of the experimental session. Specifically, we wanted to test whether the difference between the real and control conditions became smaller over time, as participants might have learned to ‘desensitize’ to the presentation of their own name, for instance. To test this, we divided trials into two sets: the first 90 trials and the last 90 trials of each participant. Then we tested these two sets separately using the same sample-by-sample linear mixed effects analysis as described above to check the effect of the critical names. In addition, in a separate sample-by-sample analysis similar to the one described above, we included both trial number and condition as fixed effects (and corresponding by-participant random effects). This analysis allowed us to check whether trial number (testing time) robustly modulates the differences between the three critical name conditions. We also considered an effect reliable if *p* < .05 for at least 200 ms (5 samples).

To further check whether our RSVP task in combination with pupillometry is useful as a tool for concealed-information detection, we wanted to establish whether the effects that were measured at the group level (see above) could also be measured reliably at the individual level.

To do this, we conducted a leave-one-out analysis both on all trials and on the first and second halves separately. For each participant separately, we first determined the peak indices (time points) at which the pupil-size difference between the real and control condition, as well as the fake and control condition, was largest for all other 30 participants. Then we conducted t-tests separately for the pupil-size difference between the real and control condition, as well as between the fake and control condition, at these corresponding peak indices. The general logic behind this approach is that we use the data from all-but-one participants as a ‘temporal localizer’ to determine the optimal time point to test for a given participant. If pupil size for the fake condition is significantly different from the control condition for each participant, then this would mean that our approach is able to detect the (attentional) reaction to task-relevant target names, even for individual participants. Similarly, if pupil size for the real condition is significantly different from the control condition for each participant, then this would mean that our approach is able to detect the presence of concealed information, even for individual participants. Given the relatively small effect sizes that are commonly observed in psychological research, and given that neither our task parameters nor analysis approach have yet been explored systematically in the present context, this is a lot to ask of our data. Nevertheless, with an eye towards practical applications, the individual-participant reliability is important to assess.

### 2.3 Results

#### 2.3.1 Task performance

Participants responded to questions 1 and 2 with an accuracy of 89.1% and 95.4% respectively. Thus, participants were well able to detect their fake names in the RSVP sequences, and to keep their attention on the stimuli throughout each trial.

#### 2.3.2 Pupil data

For the group-level analysis, the sample-by-sample linear mixed effects analysis on the full trials showed that pupil size in the fake condition was significantly larger than in the control condition from about 430ms to 2200ms (illustrated in the red straight line in Panel a). The pupil size in the real condition was significantly larger than in the control condition from about 400ms to 1000ms (illustrated in the green straight line in Panel a).

Looking at Figure 3b and 3c, in the first 90 trials, the sample-by-sample linear mixed effects analysis showed that pupil size in the fake condition was significantly larger than in the control condition from about 430ms to 2200ms (illustrated by the red straight line in Panel b). Pupil size in the real condition was significantly larger than in the control condition from about 430ms to 1250ms (illustrated by the green straight line in Panel b). In the second 90 trials, only in the fake condition was pupil size statistically significantly larger than in the control condition, from about 460ms to 2200ms (illustrated by the red straight line in Panel c). But there was no longer a reliable difference between the real and control conditions.

**Figure 3.**
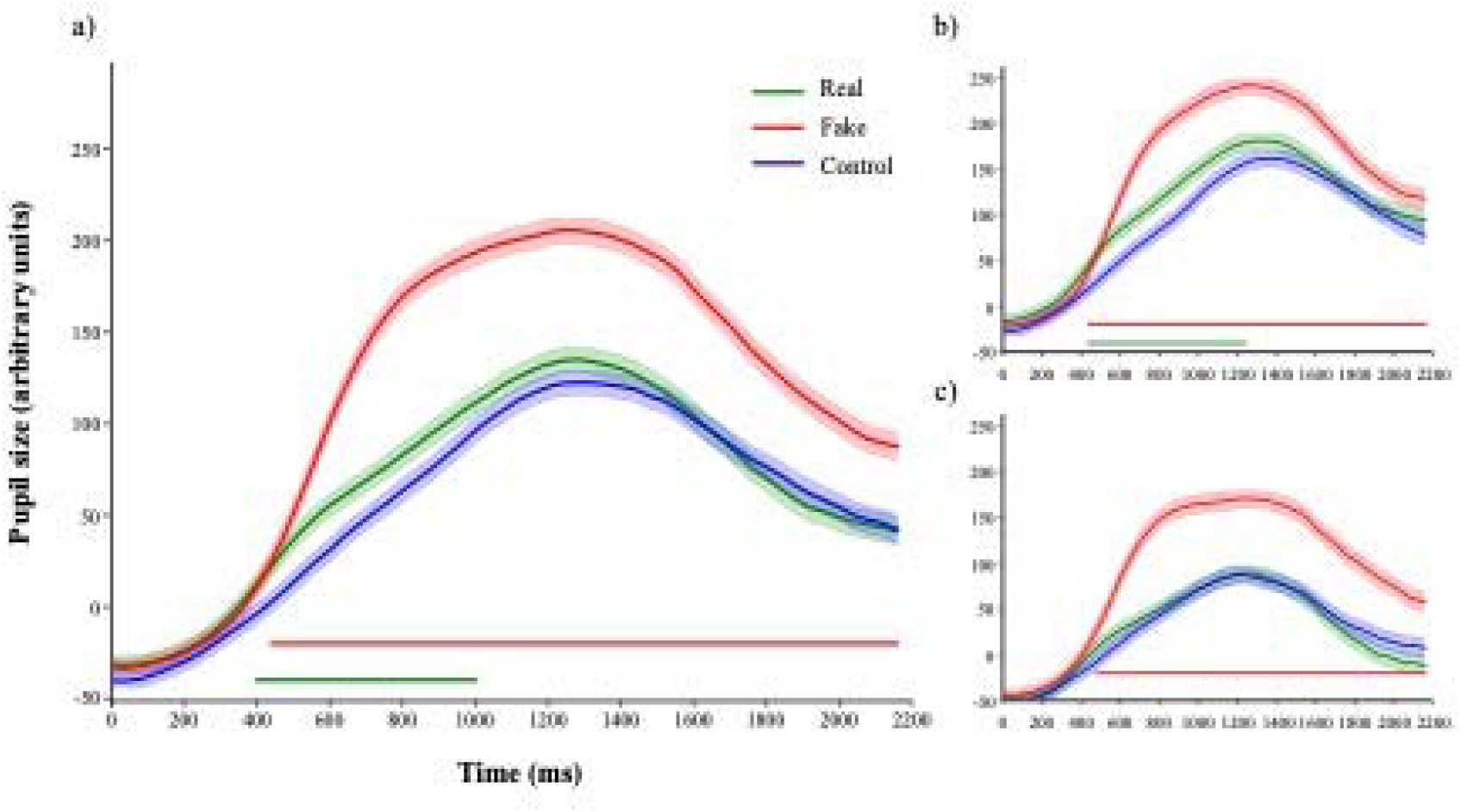
Average pupil traces. Pupil size average (N=31) in arbitrary units is shown over time (ms), in the fake condition (response to the assigned, task-relevant fake name; red line), the real condition (response to the participant’s real name; green line), and the control condition (irrelevant name; blue line) on all trials (**a**), the first (**b**) and second (**c**) half of the experiment.

However, when we tested the learning effect more rigorously as an interaction between trial number and condition, we did not find a reliable interaction (p value didn’t reach the criterion of p < .05 for at least 200ms) at any point in time. We only found that pupil size decreased over time (p < .05 for more than 200ms), possibly as an effect of increasing fatigue. Taken together, the results suggest that pupil size is sensitive to concealed identity information from the very start of the testing session.

In the individual-level leave one out analyses, although the vast majority (24 of 31) of participants showed a qualitative effect in the predicted direction, only six participants out of 31 showed a significant pupil-size difference between the real and control conditions as shown in Figure 4, Panel a. In contrast, 22 participants out of 31 showed a significant pupil-size difference between the fake and control conditions (Panel b).

**Figure 4.**
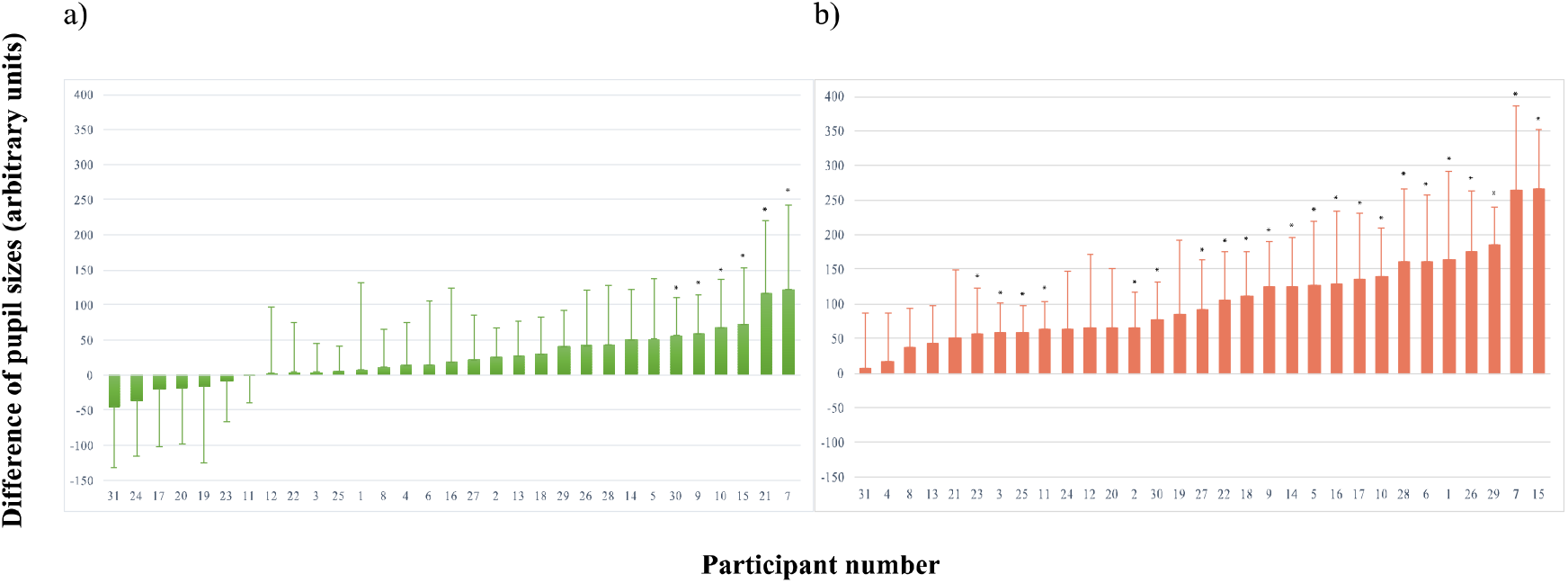
Individual effect sizes. The difference between pupil sizes for the real and control conditions of each participant (N=31, sorted by effect size) on all trials is shown in Panel **a**. The difference between pupil size for fake and control of each participant (N=31) on all trials is shown in Panel **b**. Significant differences are marked by *. The error bars indicate the individual 95% confidence intervals.

In the individual-level leave one out analyses on both half sets (90 trials each), two participants out of 31 showed a significant pupil-size difference between the real and control conditions in the first half, compared to one participant in the second half. 20 participants out of 31 showed a significant pupil-size difference between the fake and control conditions in the first half compared to 16 participants in the second half.

### 2.4 Discussion

It seems that the real names (task-irrelevant stimuli) elicited a detectable pupil reaction at the group level; fake names (the task-relevant stimuli) had a qualitatively similar but much stronger effect. The real-name effect appeared at the very beginning of the testing session, indicating that pupil size is useful as a tool for the RSVP-based concealed-information detection with short sessions. However, it was not more reliable for most participants when analysed individually on the first half of the study alone, presumably due to lower statistical power.

As an alternative to the sample-by-sample linear mixed effects analysis that we used in this experiment, it might be better to focus on a time window that is defined a priori. Specifically, 600ms to 1200ms after the onset of the critical names (corresponding to 400ms to 1000 ms in Figure 3, after subtraction of the first 200 ms) seems to be the optimal time window to assess possible differences in pupil size.

## 3 Experiment 2

The first purpose of Experiment 2 is to replicate and extend the results of Experiment 1 with a preregistered analysis plan. An important part of the analysis plan is to use the time window that we found to show the largest difference between the real and control conditions at the group level in Experiment 1. Experiment 2 was identical to Experiment 1, apart from some minor changes on the design to refine the study and attempt to enlarge the effect.

A preregistered analysis plan, all data and the analysis scripts are publicly accessible on the Open Science Framework (https://osf.io/9re75/). Deviations from the preregistration are described below.

### 3.1 Method

#### 3.1.1 Sample size

A bootstrap resampling power analysis was conducted on Experiment 1 for a sample-by-sample linear mixed effect analysis. Based on the data (N = 31), we randomly sampled a number of participants (with replacement) and ran a sample-by-sample linear mixed effect analysis. If the *p*-value for the difference of pupil size between real and control was less than .05 for more than 5 samples (200ms) in a row, we considered it as a hit. We repeated this procedure for different numbers of participants, 1000 times for each number, and determined the smallest number of participants for which a hit was obtained in 90% (power) of 1000 cases. This power analysis showed that a sample of 25 participants would be required.

#### 3.1.2 Participants

Twenty-seven participants took part in the experiment. Following the exclusion criteria defined in the preregistration, two participants were initially excluded because their accuracies for question 1 (Target search task) were below 80%. However, in deviation from the preregistration, we updated our analysis pathway to use a new, improved blink-reconstruction algorithm and to exclude trials based on deviant baseline pupil sizes (Mathôt & Vilotijević, 2022); after this update, one participant reached 80% for question 1 and was included again. Therefore, 26 participants were included in the final analysis. All participants were first-year undergraduate students at the University of Groningen and native Dutch speakers. They were in the age group of 17-25 years (Mean: 19.38 years). Among the participants were 20 females and 6 males, 25 were right-handed and 1 was left-handed. Requirements and preparation for participants, ethical approval, informed consent, debriefing and compensation received were identical to Experimental 1.

#### 3.1.3 Stimuli and Procedure

Experiment 2 was identical to Experiment 1, apart from five differences. The final four differences of the five were pre-registered. First, pupil size was recorded by an EyeLink 1000, which sampled at 1000Hz rather than 250Hz. Second, the name stimuli were presented in mono font instead of sans serif. This was to control the exact width of the presented names. Third, the participant’s real name was not presented during the practice session in order to avoid adaptation already during practice. Fourth, participants were instructed at the beginning of each block that their real name would be present in several of the streams, but that they should ignore it. This was to enhance the saliency of their real name for the participants. Fifth, a memory task was added to the end of the experiment to see which names participants remembered from the task.

During the memory task, participants were first asked to freely recall up to 5 most frequent names that they had seen during the experiment. Then, a recognition test was run, in which the critical names (real, fake and control), some noncritical names (randomly selected distractors), and three baseline names (names that had not been presented at all) were presented one-by-one. Participants rated how often they had seen the name in the experiment by giving each name a score from 1 to 5 (‘didn’t see it at all’ to ‘saw it very often’). This was done to assess the degree to which these stimuli were noticed overall.

### 3.2 Data processing and Analysis

As mentioned above, two additional data-preprocessing steps were taken that were not part of the preregistered analysis plan. These steps were based on a set of guidelines that were recently published (Mathôt & Vilotijević, 2022), and because they significantly improved data quality we felt justified in diverging from the preregistration on this point. First, we used an updated blink-reconstruction algorithm, which more effectively reduced variability due to eye blinks. Second, we removed trials where baseline pupil size deviated more than two standard deviations from the participant’s mean baseline pupil size.

We also retrospectively applied these data-preprocessing steps to Experiment 1, which also resulted in a change in the time window of interest from 500ms - 1600ms (as preregistered on open science framework) to 600ms - 1200ms.

Experiment 2 was identical to Experiment 1 in data processing and analysis, apart from four differences. First, we analyzed the rate of recall and recognition of the critical (real, fake and control), noncritical (distractors), and baseline (unpresented) names in the recall and recognition memory test for each participant. Participants were expected to recall and recognize the critical names more often than the noncritical and baseline names.

Second, we added two exclusion criteria. Participants were excluded from the pupillometric analysis and replaced by new participants if (1) less than 80% of their responses to either the question 1 (did you see your (fake) name?) or question 2 (did you see ---------- or =======?) were correct; or (2) they reported having seen more noncritical or baseline names than critical names during the memory task; or (3) they reported not having seen any critical name during the memory task. None of the participants met any of these criteria.

Moreover, on each trial, 3250 samples of pupil-size data were obtained, given that the EyeLink eye tracker sampled at 1000 Hz and a whole RSVP trial was 3250ms from the start of a name sequence to the end of the following fixation. We down-sampled the signal to 100 Hz, leaving 325 samples per trial for each participant.

Additionally, and as described above, we adopted a new analysis time window, 600ms to 1200ms after the onset of critical names, where the biggest difference between real and control was found in Experiment 1 after additional data preprocessing. The average pupil size during this time window was calculated for each trial. Then, we ran a linear mixed effect analysis for this average. Our criteria and prediction for this analysis was the same as the one for the sample-by-sample linear mixed effect analysis.

### 3.3 Results

#### 3.3.1 Task performance

Participants responded to questions 1 and 2 with an accuracy of 91% and 96.8% respectively. All participants reported having seen more critical names than noncritical or baseline names.

#### 3.3.2 Pupil data

In the group-level analysis, the sample-by-sample linear mixed effects analysis on all trials showed that pupil size in the fake condition was significantly larger than in the control condition from about 400ms to 2200ms (illustrated by the red straight line in Panel a). Pupil size in the real condition was significantly larger than in the control condition from about 420ms to 760ms (illustrated by the green straight line in Panel a).

The linear mixed effect analysis on the time window from 600ms to 1200ms after the onset of the critical names (corresponding to 400ms to 1000ms in Figure 5, after subtracting the first 200ms of deleted data) showed a significant effect both for the real condition (*t* = 2.135, *p* = .043), as well as for the fake condition (*t* = 8.158, *p* < .001).

**Figure 5.**
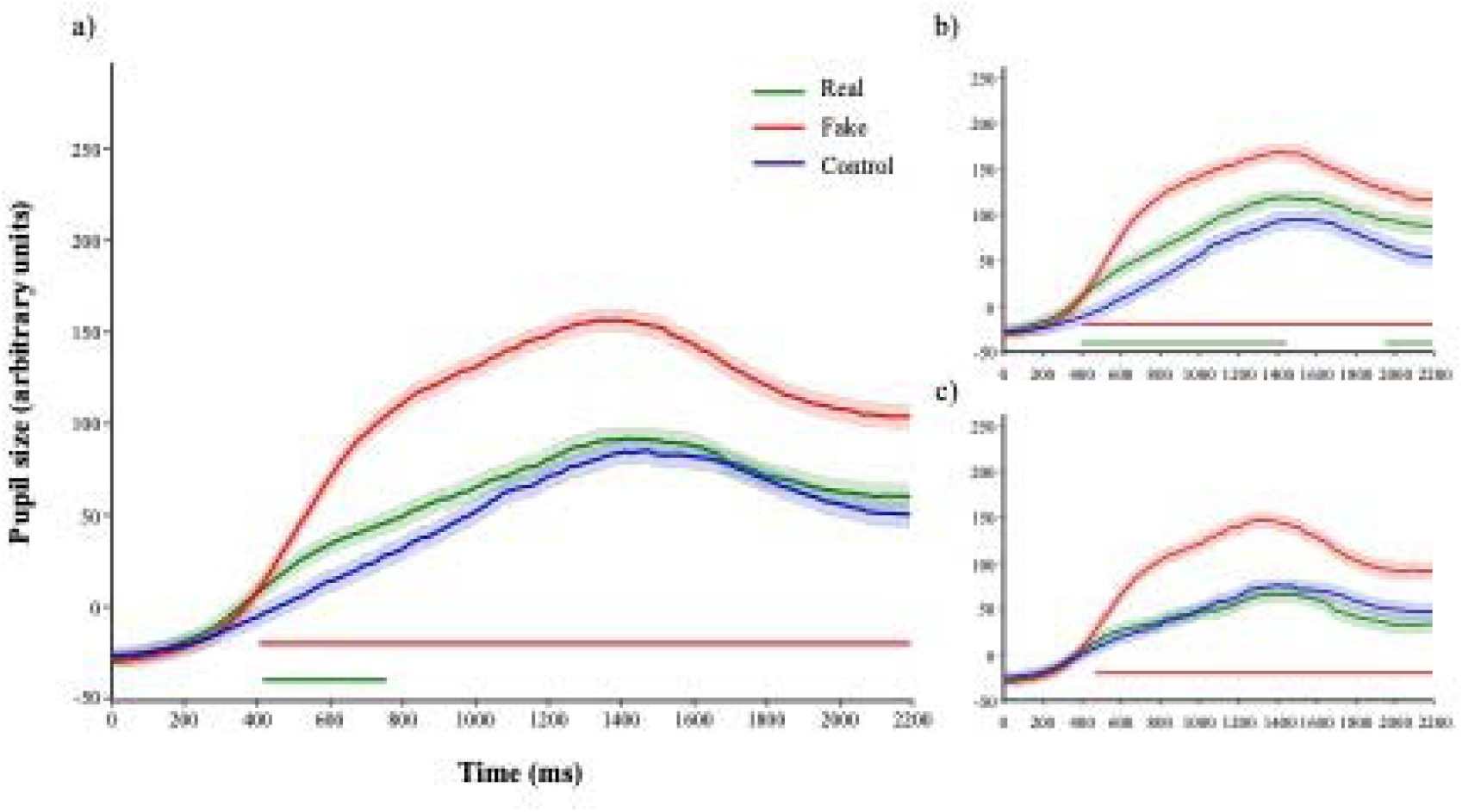
Average pupil traces. Pupil size in arbitrary units (N=26) is shown over time (ms), in the fake condition (response to the assigned, task-relevant fake name; red line), the real condition (response to the participant’s real name; green line), and the control condition (irrelevant name; blue line) in all 180 trials (**a**), and in the first (**b**) and second (**c**) half of the experiment.

Looking at Figure 5b and 5c, in the first half set of trials, the sample-by-sample linear mixed effects analysis showed that pupil size in the fake condition was significantly larger than in the control condition from about 400ms to 2200ms (illustrated by the red straight line in Panel b). Pupil size in the real condition was significantly larger than in the control condition from about 400ms to 1460ms and again from 1960ms to 2200ms (illustrated by the green straight line in Panel b). In the second half set of trials, only in the fake condition was pupil size significantly larger than in the control condition, from about 460ms to 2200ms (illustrated by the red straight line in Panel c). But there was no longer a reliable difference between the real and control conditions.

However, when we tested the learning effect more rigorously as an interaction between trial number and condition, we did not find a reliable interaction (p value didn’t reach the criterion of p < .05 for at least 200ms) at any point in time. We only found that pupil size decreased over time (p < .05 for more than 200ms), possibly as an effect of increasing fatigue.

In the individual-level leave one out analyses on all trials, which is shown in Figure 6, although the vast majority (18 of 26) of participants showed a qualitative effect in the predicted direction, only three participants showed a significant pupil-size difference between the real and control conditions (Panel a). In contrast, 17 participants showed a significant pupil-size difference between the fake and control conditions (Panel b).

**Figure 6.**
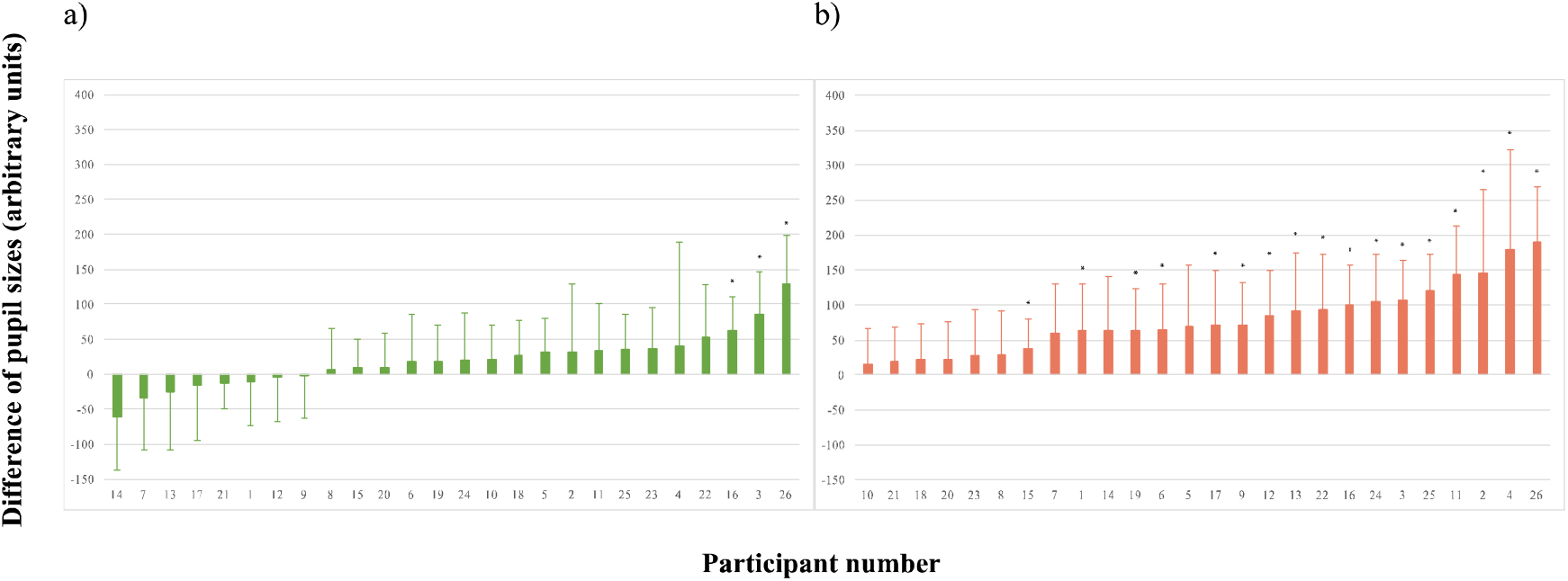
Individual effect sizes. The difference between pupil sizes for the real and control conditions of each participant (N=26, sorted by effect size) is shown in Panel **a**. The difference between pupil size for fake and control of each participant (N=26) is shown in Panel **b**. Significant differences are marked by *. The error bars indicate the individual 95% confidence intervals.

In the individual-level leave one out analyses on the first and the second halves of the experiment separately (90 trials each), four participants out of 26 showed a significant pupilsize difference between the real and control conditions on the first half compared to five participants on the second half. 13 participants showed a significant pupil-size difference between the fake and control conditions on the first half compared to 11 participants on the second half.

### 3.4 Discussion

Overall, Experiment 2 replicated the findings of Experiment 1. The real names (task-irrelevant stimuli) elicited a detectable pupil reaction in RSVP at the group level across the whole experimental session. As in Experiment 1, the real-name effect seemed most pronounced in the first half of the experiment, suggesting that long testing sessions are not necessary (and perhaps even detrimental). The time window from 600ms to 1200ms after the onset of the critical names seemed to be suitable for detecting concealed real names by pupillometry in RSVP at the group level, because in both experiments the effect emerged roughly in that period. At the individual level, the real-name was again not statistically significant for most participants.

## 4 General discussion

As shown empirically in two studies, pupil size in a rapid serial visual presentation (RSVP) task can provide valuable information regarding concealed identity information. More specifically, we observed pupil dilation in response to both participants’ ‘fake’ and real names, even though participants were instructed to respond only to the ‘fake’ name that they had selected at the start of the experiment.

The ‘fake’ name effect we found supports previous findings that pupil dilation reflects cognitive control and attention, which is required for responding to relevant information, while inhibiting irrelevant information (Cohen et al., 2015; Querino et al., 2015; Rondeel et al., 2015; van der Wel & van Steenbergen, 2018). More importantly, the ‘real’ name effect supports the finding from Gilzenrat et al. (2010)’s study that pupils also dilate in response to task-irrelevant stimuli, as long as they are salient, such as the participants’ real names in the current task. In our current study, we observed both of these two different pupil responses within one task.

Our findings are similar to the results Bowman et al. (2014) found in their EEG study. In their study, participants’ real names triggered a significant P3 component of the event-related potential, compared to control names at the group level, while they tried to search for ‘fake names’ and ignore ‘real names’. In this study, we similarly found that concealed identity information evoked a detectable pupillary response. Thus, as a cheaper, more available, less technically complex, and non-invasive measure, pupillometry is effective in RSVP for concealed identity information detection on the group level. Thereby, pupillometry can be considered as a potentially promising measure to be used widely in practice with an RSVP concealed information detector.

The pupil effect we observed also corresponds to the previous findings that the P3 component and pupil dilation both reflect the activation of phasic responses in the locus coeruleus-norepinephrine (LC-NE) system (Koss, 1986; Murphy et al., 2011; Nieuwenhuis et al., 2005, 2011; Samuels & Szabadi, 2008). Thus, it seems fair to assume that cognitive processes and attention allocation caused by the real names activated the LC-NE system, which is followed by (the P3 component and) dilated pupil size.

We furthermore observed a second practical advantage in our studies. The real-name effect on pupil size was found already at the start of the experiment and it only seemed to decline over time. Even though no evidence for a strong time-on-task effect was found, this suggests that we might be able to avoid extensive measuring sessions, which may actually reduce the effect of interest. To further benefit from the early onset of the present difference between real and control names, it may be advisable to keep real names only in the testing sessions and to not include them in the practice trials.

The current outcomes also indicated that, while most participants showed the desired effect qualitatively, it was not statistically significant for most participants when analyzed individually. This contrasts with the success rate of the EEG-based RSVP study of Bowman and colleagues (2013), in which the authors were able to increase the individual success rate considerably by using Fisher’s method to combine data from three electrodes (Fz, Cz and Pz). They succeeded in obtaining a significant p-value for the difference between the real and the control conditions for each of their 15 participants. Moreover, in the CIT study by Lubow and Fein (1996), as already mentioned in the Introduction, although participants were classified as guilty or innocent based solely on pupil size, as in our study, guilty participants were detected with 50-70% accuracy. One possibly important difference in that study compared to ours is that the authors used four different critical items taken from four categorical sets: colors, faces, signatures, and sums of money, rather than a single item.

Compared to the results of Bowman and colleagues (2013) and Lubow and Fein (1996), pupil data in the current RSVP task with the current analysis approach does not yet seem to provide the same level of sensitivity. It will be an interesting avenue for future research to attempt to boost sensitivity by combining pupil size with other measures (Seymour et al., 2013). Measures of (micro-) saccades, blink-rate, fixation duration, or even other physiological indicators, such as heart rate, may be suitable candidates. Alternatively, more cost-effective and user-friendly “dry” (i.e., without using gel as a conductor) EEG systems may provide valuable additional information, especially when combined with pupil size in the current paradigm. Using multiple critical items, as in the design of Lubow and Fein (1996), may further boost sensitivity as compared to the current approach of having only a single critical item, which may lead to habituation. Finally, using machine-learning techniques to detect subtle differences in responses to concealed items may prove more sensitive than our current analysis approach.

In conclusion, we have shown that pupil size is sensitive to the presence of concealed identity information (the participants’ own name) in an RSVP task. This implies that pupil size is a promising measure for detecting concealed information. However, further refinement is required to improve the task’s sensitivity so that concealed identity information can be detected reliably even at the level of individual participants, and we have offered several suggestions for follow-up.

## Author Notes

(1) Chen was supported by the Chinese Scholarship Council, grant 202007720091.

